# E-TAN, a technology-enhanced platform with tangible objects for the assessment of visual neglect: a multiple single-case study

**DOI:** 10.1101/849505

**Authors:** Antonio Cerrato, Daniela Pacella, Francesco Palumbo, Diane Beauvais, Michela Ponticorvo, Orazio Miglino, Paolo Bartolomeo

## Abstract

Visual neglect is a frequent and disabling consequence of right brain damage. Traditional paper-and-pencil tests of neglect have limitations in sensitivity and ecological validity. The Baking Tray Task (BTT), instead, approaches real-life situations, because it requires participants to place 16 physical objects on a board. The number of objects placed on the left and right portions of the board provides a clinical index of visual neglect. Here we present E-TAN, a technology-enhanced platform which allows patients to perform an enhanced version of the BTT (E-BTT). This platform automatically determines the object locations on the board, and also records the sequence and timing of their placement. We used E-BTT to test 9 patients with right hemisphere damage, and compared their performance with that obtained by 115 healthy participants. To this end, we developed a new method of analysis of participants’ performance, based on the use of the convex hull described by the objects on the board. This measure provides an estimate of the portion of space processed by each participant, and can effectively discriminate neglect patients from patients without neglect. E-TAN allows clinicians to assess visuospatial performance by using a convenient, fast, and relatively automatized procedure, that patients can even perform at home to follow-up the effects of rehabilitation.

## 1 Introduction

Visual neglect is a disabling pattern of spatial and nonspatial deficits, which frequently results from right hemisphere damage (Bartolomeo, 2007; Cubelli, 2017). Neglect patients behave as if the left part of the world did not exist anymore, and have poor functional outcome (Bartolomeo, 2013). Diagnosis of neglect usually relies on the administration of standard paper-and-pencil tests, such as cancellation and line bisection tasks (Wilson, Cockburn & Halligan, 1987; Azouvi, Bartolomeo, Beis, Perennou, Pradat-Diehl & Rousseaux, 2006). However, the clinical and ecological validity of these paper and pencil tests has limitations. For example, it has long being known that some patients can perform in the normal range on neglect tests, and yet show clinical signs of spatial deficits, which can emerge when questioning the patient’s carers (Azouvi, Olivier, de Montety, Samuel, Louis-Dreyfus & Tesio, 2003). In still other cases, deficits can only be unveiled by computerized response time tasks (Bartolomeo, 1997), or by tasks taxing executive attention (Bartolomeo, 2000; Bonato, 2012), which presumably prevent the use of compensatory mechanisms. These patients are at risk of being considered as having fully recovered from neglect, while still suffering from spatial deficits affecting their daily routine. These deficits can engender dangerous situations (e.g., in car driving), or induce substantial levels of handicap in everyday life (e.g., when cooking or when choosing items at a supermarket).

The Baking Tray Task (BTT) (Tham & Tegnér,1996) is a promising technique to detect the behavioral consequence of visual neglect without relying on subjective questionnaires or on computerized response time tasks. Patients are requested to dispose 16 cubes on a board as equally spaced as possible, “as if they were buns on a baking tray”. In the original study (Tham & Tegnér, 1996), no healthy participant in a group of 28 placed more than two buns in excess on one side of the board as compared with the other. On the basis of this cutoff score, seven patients with right hemisphere damage, who performed in the normal range on paper-and-pencil tests, showed neglect on the BTT. Thus, the BTT proved to be a simple and yet sensitive test of neglect. Additionally, performance seems independent of demographic variables such as participants’ age (Facchin, Beschin, Pisano & Reverberi, 2016).Given its simple and user-friendly setup, the BTT can easily be used to follow up the effects of rehabilitation (Facchin, Beschin & Daini, 2017). However, the analysis of results based on the number of items situated on the left/right halves of the board remains relatively raw, and is liable to miss subtler signs of spatial deficit, such as a pathological shrinking of the explored space. Other abnormal behavioral patters that could go undetected are the tendency to start the task from the right side (Gainotti, D’Erme & Bartolomeo, 1991), whereas healthy participants’ tendency to start from the left side (Bartolomeo, D’Erme & Gainotti, 1994; Gigliotta, Malkinson, Miglino & Bartolomeo, 2017), a general slowing of performance, or a spatially disorganized placement of items on the board, without lateralized aspects. Enhancing the BTT with technological features can enable the clinician to detect these patterns of performance.

## 2 Methods

### 2.1 E-TAN: a technologically-enhanced platform to assess visuospatial cognition with tangible interfaces

A tangible user interface system consists of an integrated system of concrete objects that participants can manipulate. The E-TAN platform is a prototype derived from the enhancement of BTT, a tool devised to evaluate of spatial cognition (Cerrato & Ponticorvo, 2017; Cerrato, Ponticorvo, Bartolomeo & Miglino, 2018; Cerrato, Ponticorvo, Gigliotta, Bartolomeo & Miglino, 2019a,b; Gentile, Cerrato & Ponticorvo, 2019). The tangible physical interfaces in E-TAN are disks 4-cm in diameter. A 30 fps camera placed on top of the board at a fixed distance, and connected to a laptop computer, can detect the disks when they disposed on a predefined board/surface, thanks to ArUco Markers (Garrido-Jurado, Muñoz-Salinas, Madrid-Cuevas & Marín-Jiménez, 2014). ArUco Marker tags are often used in augmented reality systems (Cerrato, Siano & De Marco, 2018). They consist of a pattern of black and white tiles containing compressed information, similar to QR codes (see Fig. 1).

Thus, E-TAN detects the spatial coordinates of the objects arranged on the board, and offers the opportunity to standardise the data collection and to store the subjects’ performance in both local and online databases. E-TAN also allows one to record the order and timing in which the objects are placed on the board, thus permitting the analysis of the temporal sequence of subjects’ performance.

**Figure 1:**
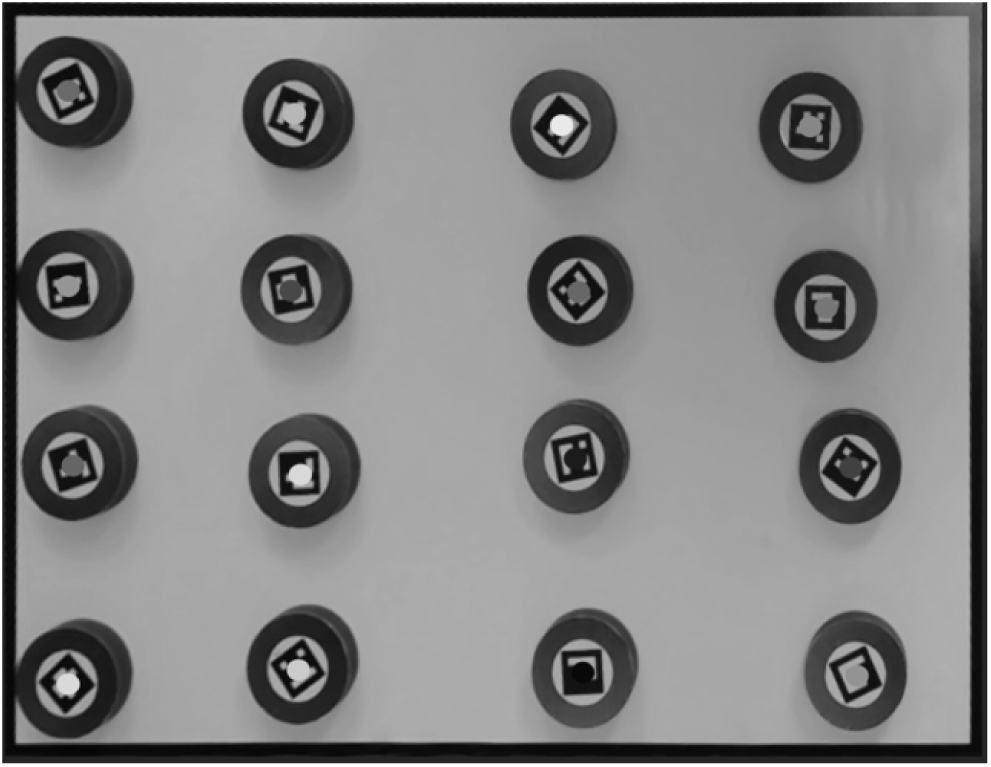
Example of E-BTT final configuration, provided by patient 6

### 2.2 Experimental procedure

The procedure was similar to that used for the traditional BTT (Tham & Tegnér, 1996; Bailey, Riddoch & Crome, 2004; Facchin et al., 2016), the only exceptions being that disks were used instead of cubes, and that the board dimensions were 60 x 45 cm. Participants were asked to dispose the disks on the board as equally spaced as possible. The administration of the task lasted about 5 minutes.

Nine right-handed patients (3 females, mean age 61.4 years, SD 13.5, range 43 - 80) were included on the basis of their having had a first stroke in the right hemisphere. The exploratory nature of the study design required no randomization or blinding/masking. Patients performed the enhanced version of the BTT (E-BTT) on the E-TAN platform at the Brain and Spine Institute (ICM), Pitié-Salpêtrière Hospital, Paris, France, following the procedures approved by the Inserm (CPP 13-41) and by the local institutional review board. No adverse events occurred. Table 1 reports demographical, clinical and neuropsychological characteristics of patients. Patients’ performance was compared with that of 115 healthy participants (61 females, mean age 27.2 years, SD 8.6, range 19-75) (Palumbo, Cerrato, Ponticorvo, Gigliotta, Bartolomeo & Miglino, 2019). Participants were tested either at the University of Naples, Italy, or at the ICM, after giving their consent according to the Declaration of Helsinki. The procedure was approved by the local institutional review boards. Performance data were collected anonymously and stored in an online database. After performing the E-BTT, some of the patients (1, 2, 4, and 6) spontaneously commented that it was an agreeable and quick task to perform, and a welcome change after so many boring paper-and-pencil tests.

**Table 1:** Demographical and clinical characteristics of patients, with their performance on the neglect battery. Gender: M, Male; F, Female. EL, educational level (1, less than 8 years of formal education; 2, 9–12 years; 3, > 12 years). Aetiology: H, haemorrhagic; I, ischemic. Asterisks indicate pathological scores compared with normative data (Azouvi et al., 2006; Gainotti et al., 1991; Rousseaux et al., 2001). Scores for landscape drawing indicate the number of omitted left-sided items. For line bisection, positive values indicate rightward deviations, negative values indicate leftward deviations.

| Patient | Gender/<br>Age/EL | Onset of<br>Illness<br>(months) | Aetiology | Visual<br>Field | Visual<br>Extinction | Bells cancellation<br>(left/right hits,<br>max = 15/15) | Landscape<br>drawing<br>(omissions) | 200-mm line<br>bisection<br>(deviation in mm) |
| --- | --- | --- | --- | --- | --- | --- | --- | --- |
| P1 | M/68/3 | 5 | H | Left inferior<br>quadrantanopia | No | 12/14 | 0.5* | -11.8* |
| P2 | F/63/3 | 12 | I | Normal | No | 9/13* | 1.5* | +1.6 |
| P3 | M/50/2 | 6 | I | Left<br>hemianopia | No | 9/13* | 2.5* | +1.6 |
| P4 | M/46/2 | 12 | H | Normal | No | 15/15 | 0 | -3.2 |
| P5 | M/55/3 | 133 | I | Normal | No | 15/14 | 0 | -5.6 |
| P6 | M/43/3 | 28 | I | Normal | No | 14/14 | 0 | -4.8 |
| P7 | M/73/1 | 123 | I | Normal | Yes | 2/11* | 1.5* | +22* |
| P8 | F/80/2 | 108 | H | Normal | Yes | 3/12* | 1.5* | +27* |
| P9 | F/75/1 | 10 | I | Normal | Yes | 3/11* | 1* | -9.2* |

### 2.3 Paper-and-pencil neglect tests

In addition to E-BTT, brain-damaged patients also performed traditional paper-and-pencil tasks from the GEREN neglect battery (Azouvi et al., 2006), including the bells cancellation task, the copy of a landscape, the overlapping figures task, and the line bisection task (see Table 1). No patient made any omissions on the overlapping figures task. For bells cancellation, we calculated the difference between omission on the left side and on the right side of the sheet, with a cutoff score of 2 (i.e., two more omissions on the left than on the right half of the sheet), as proposed by previously published guidelines (Azouvi et al., 2006). The landscape drawing consisted of a central house with two trees per side (Bartolomeo & Chokron, 1999); the total score was the sum of left-sided omissions, with each omission of a whole tree or of half of the house receiving a score of 1, and incompletely drawn items receiving 0.5. For the the line bisection task (Urbanski & Bartolomeo, 2008), patients were asked to bisect five horizontal 200-mm lines presented individually on A4 sheets of paper in landscape format (Azouvi et al., 2006). The scoring procedure consisted in measuring the deviation in mm from the true midline, averaged across the five stimuli (Azouvi et al., 2006; Rousseaux, Beis, Pradat-Diehl, Martin, Bartolomeo, Bernati, Chokron, Leclercq, Louis-Dreyfus, Marchal & others, 2001). Leftward or rightward deviations from the true center were assigned negative or positive scores, respectively. The cut-off score is −7.3 mm for leftwards deviations and +6.5 mm for rightwards deviations (Rousseaux et al., 2001).

Patients can be classified in three groups according to their performance on the paper-and-pencil test battery (Table 1). Patients 1-3 had pathological performance on two tests out of three. Patients 4-6, performed normally on all tests. Patients 7-9 obtained pathological scores on all the tests.

## 3 Results

### 3.1 Patterns of final configuration

Figure 1 displays the final disk configuration obtained by patient 6. Figure 2 shows the sequences of disk placement used by each of the 9 patients. Five patients out of 9 started to place the disks on the right side of the board, similar to the initial rightward orienting shown by right-brain damaged patients’ on different sorts of task (Gainotti et al., 1991; Jalas, Lindell, Brunila, Tenovuo & Hamalainen, 2002; Siéroff, Decaix, Chokron & Bartolomeo, 2007); six out of 9 started the configuration from the bottom side of the surface. Patient 6 started from the down-right side of the surface and then proceeded to dispose the disks vertically in four columns, occupying the whole board (see Fig. 1 and 2).

**Figure 2:**
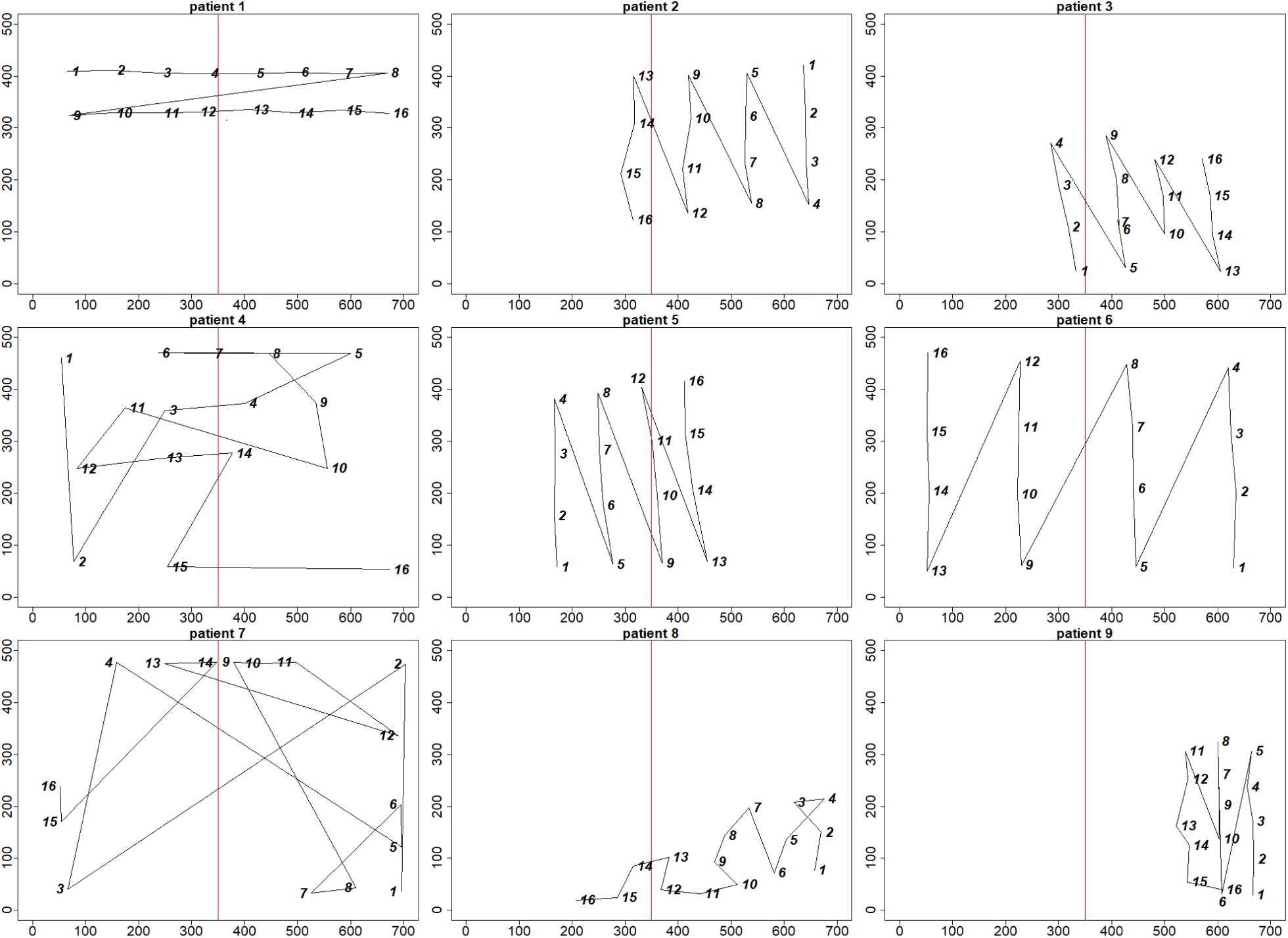
Final E-BTT conguration and sequence of disks’ placing (from 1 to 16) for each patient. Numbers represent the order of disk placement. The vertical midline of the board is represented in red.

The right-sided start might indicate a residual initial rightward bias in a patient who performs normally on paper-and-pencil tests (Bartolomeo,1997,2000; Bonato, 2012). Patients 2, 3 and 5 showed a similar sequence pattern as patient 6, but they grouped all the disks in a smaller area. In patients 2 and 3 the shrunken area is displaced rightwards, as typical of left neglect. Also the configurations of patient 8 and 9 clearly indicate severe neglect behaviors and impaired utilization of the available space. Patient 4 showed instead a generally disorganized placing strategy, without lateralized aspects.

Patient 7 paced the first 4 disks in the 4 corners of the surface (albeit without actually reaching the left upper corner), then placed the remaining 12 disks along the border of the surface, using the right-sided board limit as a guide. The final configuration remained, however, substantially asymmetrical, with only 4 disks placed in the right half of the board. Finally, patient 1 disposed the disks without any left-right asymmetry, but confined them to the upper part of the board. The scoring procedure of the traditional BTT would classify this pattern of performance as normal.

E-BTT analysis also offers the possibility of determining the localization of the first disk, which can give a clue to identify the initial attentional focus during the exploration of the surface. During neglect compensation, some patients might still initiate their exploration from the right side, and then reach the left part of the board (Bartolomeo, 1997, 2000); other patients, perhaps those who better recognize their neglect, might intentionally start their exploration from the left (Takamura, Imanishi, Osaka, Ohmatsu, Tominaga, Yamanaka, Morioka & Kawashima, 2016). E-BTT can record such shifts of exploratory preferences during patient follow-up, and thus characterize distinct behavioral patterns even in patients whose final performance reaches the normal limits. For example, in our patient sample the left or right direction of a pathological deviation on line bisection generally corresponded with the hemispace of placement of the first E-BTT object: patients who deviated leftward on line bisection also chose to place the first object on the left hemispace, and vice-versa. The the only exception was Patient 9, who deviated leftwards on line bisection but placed the first E-BTT object on the right extremity of the board. Evaluation of larger patient samples is needed to assess the generality of this relationship.

### 3.2 Advanced analysis of the exploited area

The final spatial configuration of the disks on the board, which is accurately recorded by E-TAN platform, defines the vertices of a 2D polygon. The polygon area thus represents the total space used by the patients during E-BTT performance on the left and on the right halves of the board, and can be considered as an index of the surface explored during task performance. This feature is thus likely to provide a quantitative estimate of neglect patients’ tendency to under-utilize the left portions of space. The healthy participants’ patterns of distribution of the explored spaces on each half of the board can be used as an anchor to identify pathological performance in brain-damaged patients.

Let *a* be the total covered area that is defined by the coordinates of the objects lying on the edge of the convex polygon containing the 16 disks; the Monte Carlo integration algorithm (Press & Farrar, 1990) provides an accurate estimation of *a*. The algorithm estimates the area as follows: *N* points are randomly generated from uniform distributions, points are plotted on the board, and the algorithm counts the points falling into the convex hull region. As the points are uniformly randomly generated, the covered area can be approximated by the expression *a* = *A* × *n* × *N* ^−*1*^, where *n* and *N* indicate the points falling into the convex hull and the total points, respectively; *A* = 700 × 500 = 350, 000 is the total area of the board measured in pixels. As the random process follows the Binomial random variable, with parameters *N* and *π*, with *π* unknown, *p* = *n* × *N* ^−1^ is the best estimator of *π* with standard error (SE) of order 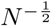, as a consequence *a* is the best estimator of the covered area, and its SE is of the same order. Analogously, the portion of *a* falling in the right/left portion of the board has been estimated and denoted with *a*_*r*_ and *a*_*l*_. Choosing *N* = 10^5^ the SE can be negligible.

A potential problem of the measure of the convex hull area is that participant may place one or a few “outlier” disks far away from the others, thus leading to an overestimation of the total area explored. To address this issue, we estimated a second left/right area measure for each participant, by removing the disks lying on the edge of the most external convex hull (such a removal technique is known as *peeling*; Eddy, 1982). Let *a*(*E*) be the area determined by the most external convex hull, and *a*(*I*) the inner part of the polygon computed without the external points (i.e. disks) of the whole area. The number of removed disks varies according to the configuration. Then, the measure assumed as proxy of the explored area is given by the mean of these two convex hull areas. Defining the following quantities *â* = (*a*(*E*) + *a*(*I*))*/*2*A, â*_*l*_ = (*a*_*l*_(*E*) + *a*_*l*_(*I*))*/A* and *â*_*r*_ = (*a*_*r*_(*E*) + *a*_*r*_(*I*))*/A*, it is possible to assume that they are determination of the *logit-normal* random variable having support in [0, 1], with unknown parameters *µ* and *σ*. The *logit-normal* is a very flexible distribution over *x* ∈ [0, 1], and by definition its *logit* function follows the Gaussian distribution. If *â* ∼ *logit* − *normal* then

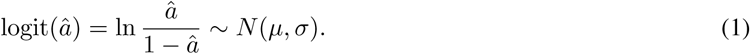

The same holds for *â*_*l*_ and *â*_*r*_.

Plots in figure 3 refer to reference population, and they show the approximation of the empirical distribution of *â, â*_*l*_, and *â*_*r*_ (in blue) with respect to the theoretical logit-normal distribution (in red). It is worth noting that the test based on the measure of discrepancy cannot reject the null hypothesis *H*_0_ : *f*_1_ ≡ *f*_2,_ where *f*_1_, *f*_2_ are the respective empirical and theoretical density functions.

**Figure 3:**
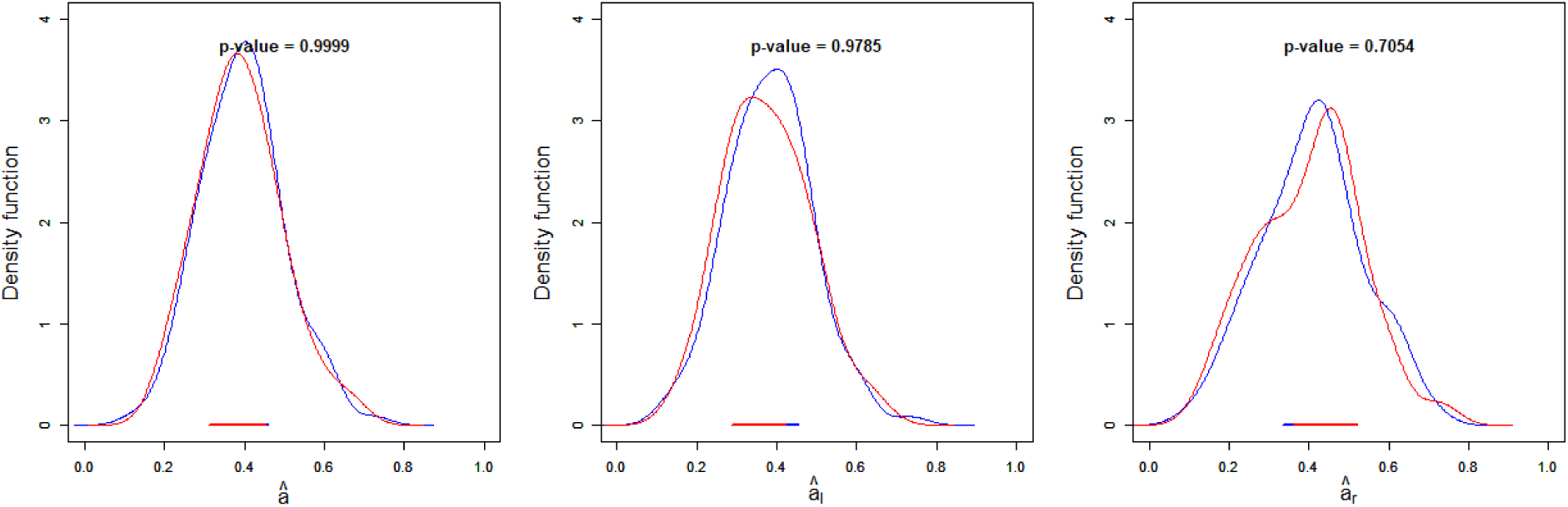
Empirical densities and theoretical logit-normal approximation with respect to the variables *â, â*_*l*_, and *â*_*r*_, from the left to the right.

The plot in Figure 4 jointly considers the standardized logit transformations of the *â*_*r*_ and *â*_*l*_ variables. The notation *lâ*_*l*_ defines the logit transformation for the left covered area while *l*â**_*r*_ defines the logit transformation for the right one. For a bivariate Gaussian the membership depends on the distance from the mean. Because the distance in more than one dimension does not have a predefined sign, a possible solution consists in considering the squared distance. It is worth to remind that the sum of *g* > 0 independent squared Gaussian random variables is distributed according to a chi-square distribution with *g* degrees of freedom (DoF). In this case the variables are not independent as the correlation index *ρ* = 68. The Mahalanobis distance breaks the correlation and the squared distances from the mean are characterised by a chi-square distribution with two DoF even if the variables are correlated. The color of the points in Figure 4 depends on membership probability to the chi-square distribution with two DoF.

**Figure 4:**
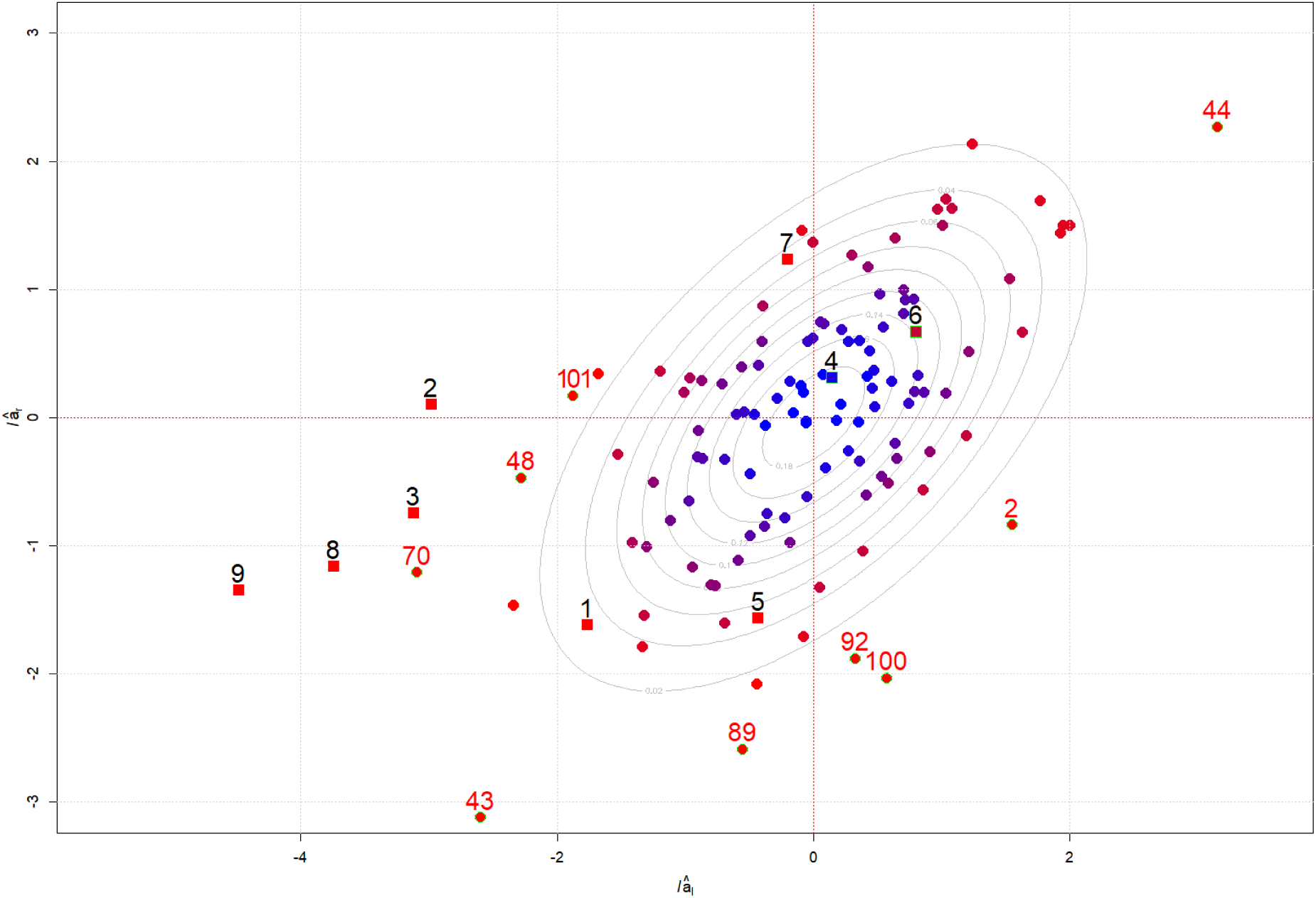
Joint plot of the standardized logit transformations of the *â*_*r*_ and *â*_*l*_ variables. In black the ID numbers of patients, in red the ID numbers of healthy subjects

Dots designate the healthy participants, squares refer to the patients with right hemisphere damage. Along with an elliptical distribution contour, points are represented according to a color gradient (from blue to red), depending on the similarity of the E-BTT configuration provided. Points with a green contour line represent those patients (4 and 6) plotted along the elliptical distribution and those healthy participants (2, 43, 44, 48, 70, 89, 92, 100, 101) that, considering their E-BTT dispositions, have a probability to belong to the “normal” group lower than .05: *P* (*z*_*l*_, *z*_*r*_ : 0, *ρ* = .68) *<* .05. It is expected that 5% of the reference population is wrongly classified as false positive. Consistent with this prediction, only 9 of the 115 healthy participants (less than 8%) showed impaired E-BTT performance. On the other hand, the performance of two of the 9 brain-damaged patients was classified as normal (patients 4 and 6 in Figure 2, who also performed paper-and-pencil tests in the normal range). Note, however, that patient 4 displayed a spatially disordered, although nonlateralized, pattern of object placement; as already mentioned, patient 6 started from the low right corner. Additional methods of analysis, e.g. focusing on spatial entropy (Batty, 1974) and sequence of placement, may increase the discrimination power of E-BTT.

Another possibility is to compute the ‘center of mass’ of the explored surface (Rorden & Karnath, 2010; Gainotti, Perri & Cappa, 2002). Patient 7 is plotted on the uppermost part of distribution, because he placed all the disks along the borders of the surface, and thus used a wider area than expected for typical neglect-related configurations. Performance of patients 1 and 5 was plotted in the inferior external part of the distribution, far from the center, because they covered a relatively small area of the board.

Table 2 compares the results of the present explored area method with the original BTT scoring (Tham & Tegnér, 1996)), and with the BTT bias, a scoring method which determined normative data for clinical use of the traditional BTT, by using the percentage ratio of the difference between right- and left-sided objects divided by the total number of placed objects (Facchin et al., 2016).

**Table 2:**
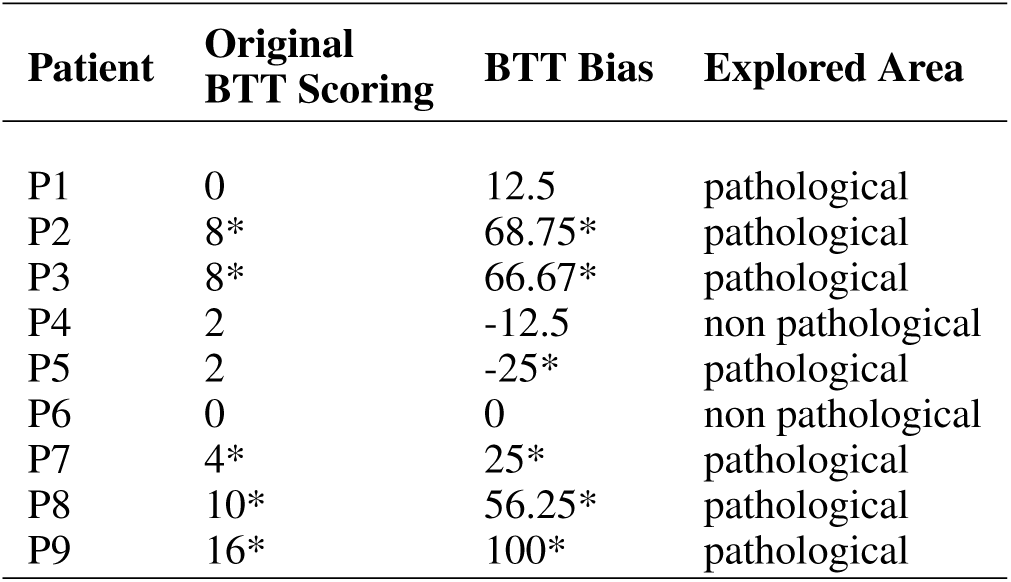
Three types of scoring (BTT, BTT Bias and Explored Area). Asterisks indicate pathological scores. For the Explored Area, the statistical procedure showed in the present paper, a dichotomic classification pathological/non pathological has been used. Original BTT scores greater than 2 indicate left neglect. BTT bias scores less than −12.5% indicate pathological leftward bias (right neglect); values greater than 18.7% indicate pathological rightward bias (left neglect).

| Patient | Original BTT Scoring | BTT Bias | Explored Area |
| --- | --- | --- | --- |
| P1 | 0 | 12.5 | pathological |
| P2 | 8* | 68.75* | pathological |
| P3 | 8* | 66.67* | pathological |
| P4 | 2 | -12.5 | non pathological |
| P5 | 2 | -25* | pathological |
| P6 | 0 | 0 | non pathological |
| P7 | 4* | 25* | pathological |
| P8 | 10* | 56.25* | pathological |
| P9 | 16* | 100* | pathological |

When applied to our patient sample, the BTT bias (Facchin et al., 2016) led to similar conclusions, except for Patient 1, whose performance was classified as normal by the BTT bias scoring method, but as pathological by our method.

## 4 Discussion

Signs of visual neglect are likely to result from the interaction of multiple deficits, with substantial variability from patient to patient (Bartolomeo,2007; Siéroff et al.,2007). Thus, it is not surprising that a battery of tests has more diagnostic sensitivity that any single test alone (Azouvi et al., 2006). However, patients often anage to achieve normal performance on paper-and-pencil tests, while still suffering from substantial levels of clinical disability. The administration of E-BTT through the E-TAN platform allows clinicians to assess visuospatial performance by using a convenient, fast, and relatively automatized procedure. The recorded data can be anonymized and sent to an online platform for automatic analysis and storage for online or offline reference.

E-BTT keeps the ecological flavor of the traditional BTT, which is close to everyday life use and manipulation of objects (Miglino & Ponticorvo, 2018; Di Fuccio, Ponticorvo, Ferrara & Miglino, 2016); however, E-BTT also offers sensitivity to subtle forms of spatial deficits, thanks to a novel, advanced statistical procedure to assess patients’ patterns of performance.

In addition, the E-BTT board requires patients to reach further in the left space than during typical paper-and-pencil tests. This characteristic opens the possibility of unveiling instances of reduced leftward hand movements, or directional hypokinesia (Bartolomeo, D’Erme, Perri & Gainotti, 1998). The E-TAN platform is easily portable and can thus be used to follow-up patients during home-based rehabilitation (Rossit, Benwell, Szymanek, Learmonth, McKernan-Ward, Corrigan, Muir, Reeves, Duncan, Birschel, Roberts, Livingstone, Jackson, Castle & Harvey, 2019).

Assessment procedures exploiting the usage of tangible objects are increasingly adopted for both diagnostic and rehabilitation purposes (Ferrara, Ponticorvo, Di Ferdinando & Miglino, Cerrato, Siano, De Marco & Ricci, 2019; Rabuffetti, Meriggi, Pagliari, Bartolomeo & Ferrarin, 2016). Previous examples of technology-enhanced assessment of spatial deficits include virtual reality mazes (Morganti & Riva, 2014), which however do not require any manipulation of tangible objects. Other examples include a technology-enhanced line cancellation task (D’Amico, Landucci & Pezzatini, 2013), and a mixed reality interface equipped with tangible objects and a virtual reality environment to evaluate and rehabilitate cognitive mistakes in daily tasks (Edmans, Gladman, Walker, Sunderland, Porter & Fraser, 2007). These systems are mainly focused on rehabilitation, and lack an in-depth performance analysis. They either employ standard evaluation procedures of the traditional line cancellation task (Albert, 1973), or need the expert evaluation of an occupational therapist. The *SIG-Blocks* interface (Lee, Jeong, Schindler & Short, 2016) makes use of tangible objects equipped with motion sensors for cognitive assessment, based on some items of the Wechsler Adult Intelligent Scale. Another technology-enhanced assessment and training tool, *MOTO tiles* (Liu, Lund & Wu, 2018), consists of interactive modular tiles similar to jigsaw pieces. The tiles need to be placed one next to the other to compose a sort of mat, with audio and visual stimuli. Also this platform, however, lacks the possibility of systematic data collection and monitoring of performance.

## 5 Conclusions and future directions

E-BTT, administered through the E-TAN platform, allows clinicians to assess patients’ patterns of performance not only on the basis of the raw count of the objects disposed in each half of the board, but also by recording the spatial coordinates of the objects. In addition, other analysis techniques will be developed to increase E-BTT’s discriminating power, based on the sequence, timing and order of the patterns arranged by participants, and on the density of the disks placed in different portions of the board. In particular, the study of the regularity of object placement can provide information on poorly known clinical conditions, such as nonlateralized spatial deficits after brain damage. Because of its fast, easy and user-friendly procedure, the E-TAN platform seems also ideally suited for test-retest procedures in follow-up studies, and as a means to assess the efficacy of rehabilitation interventions. A commercial version of the E-TAN platform will be made available shortly. Clinicians will be able to easily develop or reproduce additional neuropsychological tasks to assess other cognitive abilities, such as numerical cognition or language (De Renzi & Faglioni, 1978).

